# Mitochondria-derived vesicles deliver antimicrobial payload to control phagosomal bacteria

**DOI:** 10.1101/277004

**Authors:** Basel H. Abuaita, Tracey L. Schultz, Mary X. O’Riordan

## Abstract

Pathogenic bacteria taken up into the macrophage phagosome are the target of many anti-microbial effector molecules. Although mitochondria-derived antimicrobial effectors such as reactive oxygen species (mROS) are reported to aid in bacterial killing, it is unclear how these effectors reach bacteria within the phagosomal lumen. To examine the crosstalk between mitochondria and phagosomes, we monitored the production and the spatial localization of mROS during methicillin-resistant *Staphylococcus aureus* (MRSA) infection. We showed here mROS, specifically hydrogen peroxide (mH_2_O_2_) can be delivered into phagosomes via infection-induced mitochondria-derived vesicles, which are generated in a Parkin-dependent manner. Accumulation of mH_2_O_2_ in phagosomes required TLR signaling and the mitochondrial superoxide dismutase, Sod2, which converts superoxide into mH_2_O_2_. These data highlight a novel mechanism by which the mitochondrial redox capacity enhances macrophage antimicrobial function by delivering mitochondria-derived effector molecules into bacteria-containing phagosomes.

## Introduction

Reactive oxygen species (ROS) play pivotal roles in signaling and defense of biological organisms. These highly reactive molecules oxidize lipids, proteins and other cellular constituents, leading to a spectrum of responses ranging from altered signaling to cell death. While ROS are constitutively generated and de-toxified during cellular metabolism, ROS levels acutely increase during cellular stress (Holmstrom and Finkel, 2014), and can be a potent weapon in the host arsenal to control invading pathogens. In cells of the innate immune system, like macrophages, ROS are primarily generated by the phagocyte NADH oxidase and mitochondrial metabolism. Upon infection, the phagocyte oxidase multi-protein complex, also referred to as NOX2, can be recruited to phagosomal membranes to generate a burst of superoxide into the phagosome lumen (Winterbourn and Kettle, 2013). However, many bacterial pathogens, like *Mycobacterium tuberculosis, Salmonella enterica* serovar Typhimurium, *Coxiella burnetti*, and *Francisella tularensis* prevent the rapid NOX2-mediated burst either by avoiding complex recruitment or by utilizing detoxification mechanisms (Celli and Zahrt, 2013; Koster et al., 2017; Mertens and Samuel, 2012; Vazquez-Torres and Fang, 2001). Mitochondrial ROS (mROS) can also contribute to bacterial killing (West et al., 2011), but how these reactive molecules reach bacteria within the macrophage phagosome is ill-defined.

ROS induction can be driven by cellular stress response pathways (Bronner et al., 2015), like the endoplasmic reticulum unfolded protein response (UPR), which is activated by the three ER sensors, PERK, ATF6 and IRE1*α* (Zeeshan et al., 2016; Zhang and Kaufman, 2004). Indeed, many studies support an integral role for cellular stress pathways in modulating innate immune responses (Muralidharan and Mandrekar, 2013). We recently showed that IRE1*α* is critical for stimulating macrophage anti-microbial function (Abuaita et al., 2015). Specifically, IRE1*α* activation resulted in sustained macrophage ROS production required for killing MRSA. Notably, IRE1*α* dependent killing was only partially NOX2-dependent, despite the well established role of NOX2 in neutrophil oxidative defenses against *Staphylococcus aureus (Rigby and DeLeo, 2012)*. Since IRE1*α* signaling induces mROS production (Tufanli et al., 2017), we hypothesized that the mechanism by which IRE1*α* activation led to MRSA killing relied on mROS generation.

Here we show that infection of macrophages by MRSA stimulates IRE1*α*-dependent production of mROS, specifically hydrogen peroxide mH_2_O_2_. Infection also triggers generation of Parkin-dependent mitochondrial-derived vesicles (MDVs), previously described as a pathway for mitochondrial quality control (Soubannier et al., 2012a). These MDVs deliver the mitochondrial peroxide-generating enzyme, Sod2, into the bacteria-containing phagosome, controlling bacterial burden. Our findings reveal a mechanism by which programmed cellular stress responses repurpose a mitochondrial quality control mechanism to enable anti-microbial defense.

## Results

### Induction of mROS by IRE1α promotes macrophage bactericidal function

We previously showed that MRSA infection stimulates macrophages to produce global ROS via the activation of the ER stress sensor IRE1*α*, which was required *in vitro* and *in vivo* for effective MRSA killing (Abuaita et al., 2015). Although the phagocyte NADPH oxidase-2 (NOX2) complex contributed somewhat to MRSA killing by macrophages, IRE1*α* stimulated robust macrophage bactericidal activity even in the absence of NOX2. Previous reports indicated that mitochondrial ROS (mROS) enhance macrophage bactericidal function (Geng et al., 2015; West et al., 2011), so we hypothesized that IRE1*α*-dependent antimicrobial activity might rely specifically on mROS. To test this hypothesis, we assessed whether mROS was induced by macrophages in response to MRSA infection. First, we validated that MitoPY1, a mitochondrially targeted probe that fluoresces in response to mH_2_O_2_ (Dickinson and Chang, 2008; Dickinson et al., 2013), was indeed mitochondrially restricted in macrophages. RAW264.7 cells were transfected with mito-mCherry and loaded with MitoPY1. MitoPY1 fluorescence intensity increased when macrophages were stimulated with exogenous H_2_O_2_, and the signal co-localized with mito-mCherry (Fig. 1A). We also quantified the difference in mean fluorescence intensity (MFI) by flow cytometry (Fig. 1B). During MRSA infection, MitoPY1 fluorescence intensity increased over time, peaking at 4h pi (Fig. 1C). To determine whether IRE1*α* was required for mH_2_O_2_ induction by MRSA infection, we generated IRE1*α*-deficient macrophages using CRISPR/Cas9 (Fig. 1D and S1). IRE1*α* deficiency suppressed the ability of macrophages to induce mH_2_O_2_ during MRSA infection when compared to non-target control (Fig. 1E). To test the requirement for mROS in killing MRSA, infected macrophages were treated with the ROS scavenger, NecroX-5, which is primarily localized to mitochondria (Kim et al., 2010; Thu et al., 2016). Infected macrophages treated with NecroX-5 exhibited lower MitoPY1 fluorescence than control-treated cells (Fig. 1F), and decreased capacity to kill MRSA (Fig. 1G). These data indicate that IRE1*α* is critical for infection-induced mH_2_O_2_, and establishes a role for mROS in macrophage bactericidal function against MRSA.

**Figure 1.**
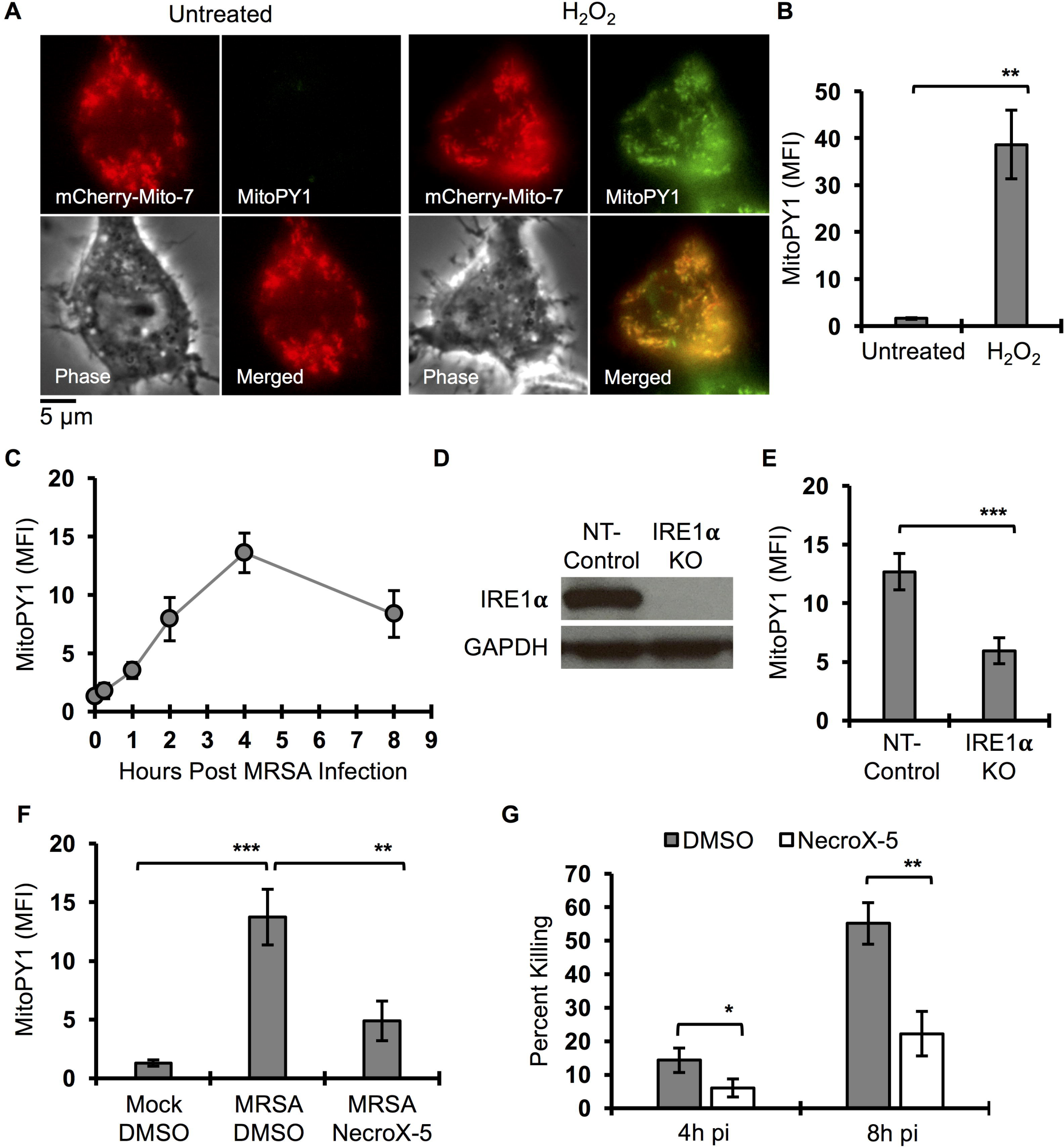
MRSA infection stimulates bactericidal mH_2_O_2_ via IRE1α. (A) Representative live fluorescent images of macrophages transfected with mCherry-Mito-7 encoded plasmid, pulsed with MitoPY1 for 1h and chased with H_2_O_2_ (100 µM) or left untreated. Images were acquired by Olympus IX-70 inverted microscope and analyzed by MetaMorph for Olympus imaging software. (B) Flow Cytometry mean fluorescent intensity (MFI) of macrophages after pulse with MitoPY1 for 1h and chased with H_2_O_2_ or left untreated. (C) Time course measurement of MFI of macrophages when pulsed with MitoPY1 and an infected with MRSA and monitored over time via flow cytometry. (D) Immunoblots of IRE1*α* and GAPHD from cell lysates from NT-control and IRE1*α* KO macrophages. (E) MFI of IRE1*α* KO macrophages and NT-control after labeling with MitoPY1 followed by MRSA infection and analyzed after 4h. (F) MFI of macrophages labeled with MitoPY1 for 1h and infected with MRSA for 4h in the presence and absence of mROS scavenger NecroX-5. (G) Percent of intracellular MRSA killing by macrophages in the presence and absence of NecroX-5. Percent killing was calculated by the following formula [1 - (CFU _indicated time_ points / CFU_1h pi_)] × 100, which represents the percent difference in CFU at indicated time point relative to 1h pi. MFI of MitoPY1 quantification was determined by using FlowJo software, representing a geometric mean. MFI of each condition was subtracted from MFI obtained from unstained cells. Graphs are presented as mean of n≥3 independent experiments +/− SD. pValue: * < 0.05, **< 0.01 and ***< 0.001.

### Mitochondrial peroxide accumulation in phagosomes is TLR-dependent

We reasoned that mH_2_O_2_ could contribute to bactericidal function indirectly by signaling and/or by direct delivery to the phagosome. If direct delivery, we might expect to see mH_2_O_2_ accumulate in the phagosome. To monitor mH_2_O_2_ spatial localization during infection, we imaged live cells stimulated with viable MRSA, killed MRSA or latex beads. Macrophages were pulsed with MitoPY1 and chased 4h post-phagocytosis (Fig. 2A). Hydrogen peroxide increased within the mitochondrial network during infection with live or fixed MRSA, but not with beads. We also observed smaller MitoPy1^+^ puncta throughout the cell. Notably, mH_2_O_2_ accumulated in MRSA-containing phagosomes (Fig. 2A and 2B). To visualize the dynamic distribution of mH_2_O_2_ during infection, we performed time-lapse imaging of infected macrophages pre-loaded with MitoPY1 (Movie S1). By 10 min pi, the MitoPY1 signal within the mitochondrial network had increased. MitoPy1^+^ puncta first associated with the bacterial phagosome at approximately 50 min pi, followed by accumulation of probe within the bacteria-containing phagosome (Fig. 2C and Movie S1). These data suggest that mitochondrially-derived hydrogen peroxide accumulates within phagosomes, and reveal the possibility that mH_2_O_2_ may contribute to macrophage bactericidal effector function through a direct delivery mechanism.

**Figure 2.**
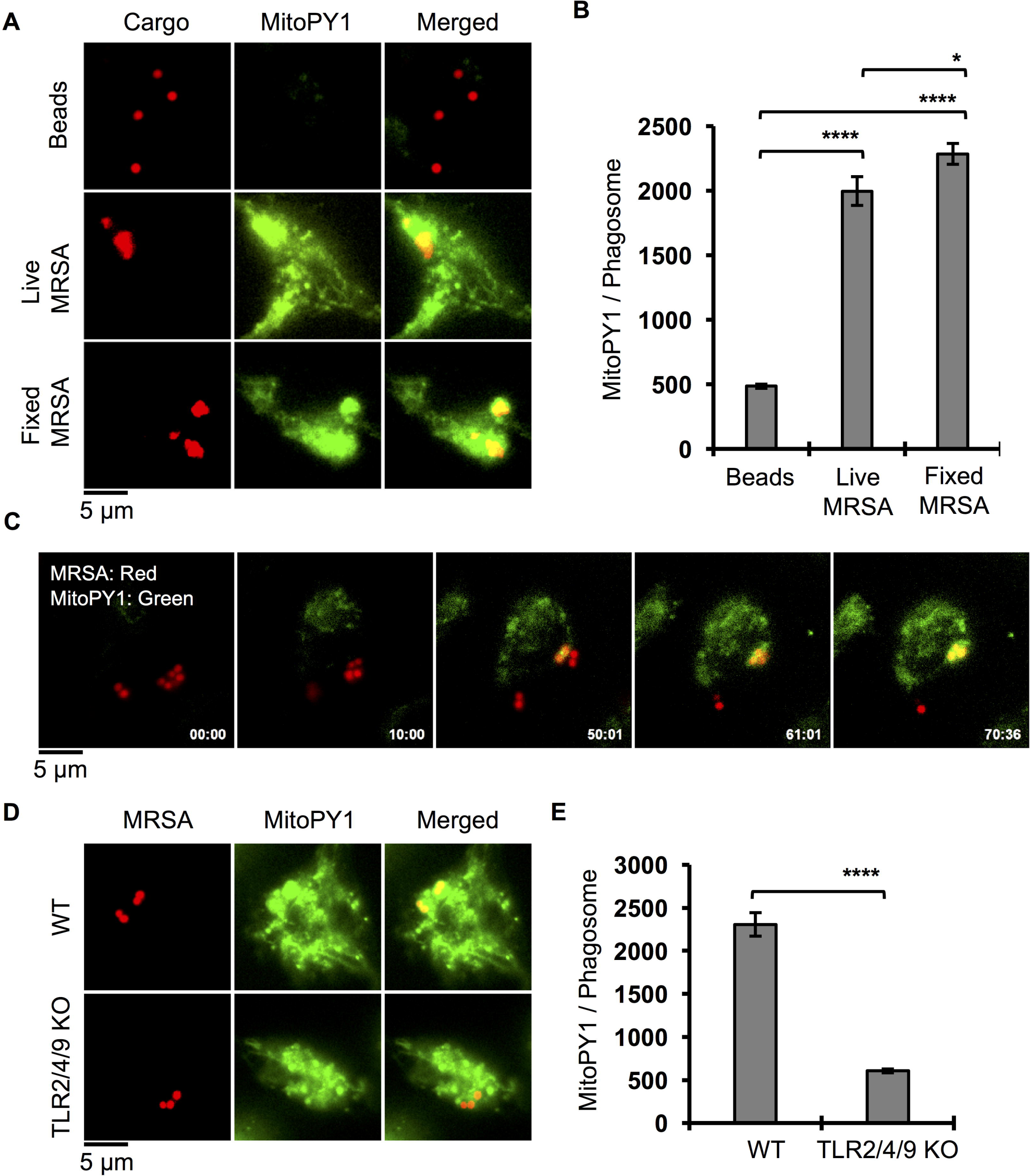
TLR signaling controls mH_2_O_2_ accumulation in the bacterial phagosome. (A) Representative fluorescent microscopy images of macrophages when pulsed with MitoPY1 (green) for 1h and chased with beads (red) or live and dead MRSA-mcherry (red) infection for 4h. Images were acquired by an Olympus IX-70 inverted live-cell fluorescence microscope and analyzed by MetaMorph imaging software. (B) Quantification of mean fluorescent intensity (MFI) of MitoPY1 associated with macrophage phagosomes using imageJ software. Phagosomes were defined by the area in the cell where red fluorescent beads or MRSA are localized. (C) Time lapse imaging of macrophages pulsed with MitoPY1 for 1h (green) and then infected with MRSA-mCherry (red). Time clock Minutes : Seconds. (D) Live microscopy images of wild type (WT) and TLR 2/4/9 deficient (TLR2/4/9 KO) macrophages pulsed with MitoPY1 (green) for 1h and then infected with MRSA-mCherry (red) for 4h. (E) Quantification of MFI of MitoPY1 associated with phagosomes from WT and TLR2/4/9 deficient macrophages using similar criteria as in panel B. Graphs represent averages of MitoPY1 MFI from at least 305 phagosomes pooled from at least three independent experiments. Error bars represent standard error of the mean (SEM). pValue: * < 0.05 and **** < 0.0001.

Mitochondrial ROS induction in macrophages occurs when cells internalize beads conjugated to Toll-like receptor (TLR) ligands (TLR2 and TLR4), but not beads alone (West et al., 2011). To test whether TLR signaling was required for mH_2_O_2_ generation and accumulation within MRSA-containing phagosomes, we measured MitoPY1 fluorescence intensity in wild-type (WT) and TLR2/4/9-deficient bone marrow-derived macrophages (BMDMs) during MRSA infection. We observed increased overall mH_2_O_2_ in both WT and TLR2/4/9 deficient BMDMs during MRSA infection, indicating that mH_2_O_2_ induction is largely independent of TLR2/4/9 signaling (Fig. S2A). However, TLR2/4/9-deficient macrophages failed to kill MRSA (Fig. S2B) and MitoPY1 accumulation in their MRSA-containing phagosomes was decreased compared to WT macrophages (Fig. 2D and 2E), indicating that TLR signaling controls mH_2_O_2_ delivery to or accumulation within bacteria-containing phagosomes.

### Bacterial infection triggers Parkin-dependent generation of mitochondrial-derived vesicles

Recent studies have revealed that mitochondrial constituents can be selectively transported to other intracellular compartments by small mitochondrial-derived vesicles (MDVs) (McLelland et al., 2014; Soubannier et al., 2012a). MDVs form in an early dynamin related protein 1 (Drp1)-independent response to increased oxidative stress, and are positive for mitochondrial markers, like Tom20. MDVs may function as a quality control mechanism by delivering damaged mitochondrial components to the endolysosomal pathway (Sugiura et al., 2014). A subset of MDVs associate with peroxisomes, and may represent a mechanism of communication between these two organelles (Neuspiel et al., 2008). Since infection increases mH_2_O_2_, we reasoned that could trigger generation of MDVs, a potential mechanism to deliver antimicrobial content from the mitochondria to the phagosome. To test this hypothesis, we first assessed whether MDVs are induced by MRSA infection. Macrophages were infected with MRSA, and subjected to immunofluorescence analysis by high-resolution confocal microscopy using a Tom20-specific antibody. MRSA infection stimulated an increase of small Tom20^+^ particles, here referred to as MDVs, compared to bead-containing macrophages (Fig. 3A and 3B, Fig. S3). The Parkinson’s Disease associated protein, Parkin, regulates biogenesis of a subset of MDVs (McLelland et al., 2014). We therefore tested the requirement for Parkin in generating infection-induced MDVs. WT and Parkin-deficient (*Park2*^*-/-*^) BMDM were infected with MRSA, and accumulation of Tom20^+^ MDVs visualized by confocal microscopy. MRSA-infected *Park2*^*-/-*^ BMDM had significantly lower numbers of Tom20^+^ MDVs compared to WT BMDM (Fig. 3C and 3D). Collectively, these results show that MRSA infection induces formation of Tom20^+^ MDVs through a Parkin-dependent mechanism.

**Figure 3.**
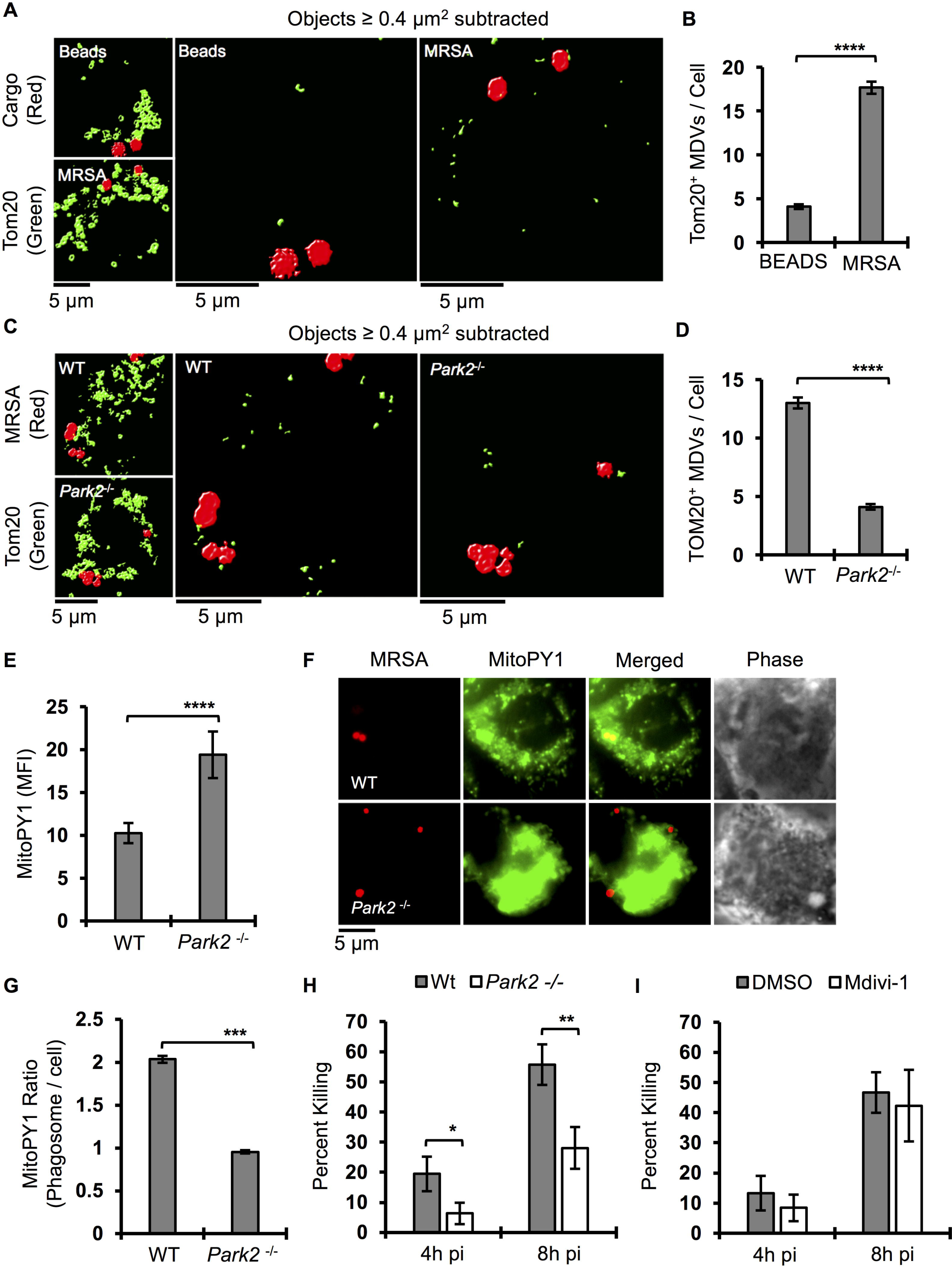
Infection induced Parkin-dependent MDVs contribute to MRSA killing. (A) Confocal microscopy representative images of macrophages stimulated with beads (red) or infected with MRSA (Red) for 4h and stained with Tom20 (green) antibody. Images were acquired by Leica TCS SP8 scanning confocal microscope and deconvoluted using Huygens essential software. Right panel are processed images after subtraction of Tom20 positive large objects (surface area > 0.4 µm^2^). (B) Quantification of Tom20 positive smaller objects (surface area < 0.4 µm^2^) per macrophage when stimulated with beads or infected with MRSA. Decovoluted confocal images were processed by Huygens essential software using the following criteria; 10% threshold, 10% seed and garbage of 50. Tom20 positive objects with surface area larger than 0.4 µm^2^ were filtered out and the remaining objects were enumerated per cell basis. Graphs represent means of at least 107 cells pooled from three independent experiments +/− SEM. (C) Representative confocal microscopy images of WT and Parkin deficient (*Park2*^-/-^) macrophages infected with MRSA for 4h. Images were processed using similar criteria as in panel A. (D) Quantification of Tom20 positive smaller objects (surface area < 0.4 µm^2^) from WT and Parkin deficient macrophages infected with MRSA. Confocal microscopy images processed as in panel B and presented as mean +/− SEM from at least 135 cells pooled from three independent experiments. (E) Flow cytometry mean fluorescence intensity (MFI) of WT and Parkin deficient macrophages when pulsed with MitoPY1 for 1h and infected with MRSA for 4h. Data represented as geometric mean of n≥3 independent experiments +/− SD. (F) Representative live fluorescence wide-filed microscopy images of WT and Parkin deficient macrophages pulsed with MitoPY1 (green) for 1h and infected with MRSA-mCherry (red). Images were acquired at 4h pi and processed by MetaMorph software. (G) Ratiometric measurement of MitoPY1 fluorescent intensity of phagosome relative to total cellular fluorescent intensity (MFI-phagosome/MFI-cell). Phagosomes were defined by the area of the cell where the red fluorescent MRSA-mCherry are located. (H) Percent of MRSA intracellular killing by WT and Parkin deficient macrophages was quantified by the following formula [1 - (CFU _indicated time points_ / CFU_1h pi_)] × 100, which represent the percent difference in CFU obtained at indicated time point relative to 1h pi. Data are presented as mean of n≥3 independent experiments +/− SD. (I) Percent of MRSA intracellular killing by RAW264.7 macrophages when treated with control DMSO or Dynamin-related protein 1 (Drp1) selective inhibitor (Mdivi-1, 25 µM). pValue: * < 0.05, **< 0.01, ***< 0.001 and ****< 0.0001.

### Parkin controls mH_2_O_2_ accumulation in phagosomes and promotes bactericidal effector function

Polymorphic alleles of *Park2* can mediate susceptibility to microbial infection (Al-Qahtani et al., 2016; Manzanillo et al., 2013). We hypothesized that MDVs could enhance macrophage bactericidal function by facilitating mH_2_O_2_ accumulation within phagosomes. We first measured the requirement for Parkin in mH_2_O_2_ accumulation in phagosomes. Upon infection, Parkin-deficient macrophages produced higher levels of mH_2_O_2_ than WT macrophages, indicating that Parkin is not necessary for mH_2_O_2_ induction (Fig. 3E). Notably, despite higher overall levels of mH_2_O_2_, Parkin-deficient macrophages displayed only minimal accumulation of MitoPY1 in phagosomes compared to WT macrophages (Fig. 3F and 3G). To evaluate the contribution of Parkin to macrophage bactericidal capacity against MRSA, WT and *Park2*^*-/-*^ BMDM were infected with MRSA to assess killing. Parkin-deficient macrophages were less capable of killing MRSA compared to WT macrophages (Fig. 3H). Parkin can regulate mitochondria quality control by inducing mitophagy (Matsuda et al., 2010), or by formation of MDVs (McLelland et al., 2014). To determine if the Parkin-dependent killing mechanism we observed involved mitophagy, we measured macrophage bactericidal function in the presence of Mdivi-1, a small molecule of Drp1, required for mitophagy (Narendra et al., 2008), but not MDV scission (Soubannier et al., 2012a). Macrophages treated with Mdivi-1 killed MRSA as efficiently as DMSO-treated macrophages (Fig. 3I).

These data indicate that Parkin-dependent generation of MDV enables accumulation of bactericidal mH_2_O_2_ in the macrophage phagosome.

We then tested the role of Parkin in innate immunity against MRSA using a subcutaneous infection model, where innate immune defenses are essential for bacterial clearance (Tseng et al., 2011). WT and Parkin-deficient mice were inoculated with 10^7^ MRSA subcutaneously into the shaved flank. Skin lesions were excised at 3 days pi, and bacterial burden and pro-inflammatory cytokines were measured. Lesions from Parkin-deficient mice yielded higher bacterial numbers (Fig. 4A), and increased levels of KC and IL-1*β* compared to WT mice (Fig. 4B and 4C). These data support a role for Parkin in innate immunity against MRSA infection *in vivo* and suggest that parkin mediates macrophage bactericidal function via delivering Tom20^+^ MDV into phagosomes.

**Figure 4.**
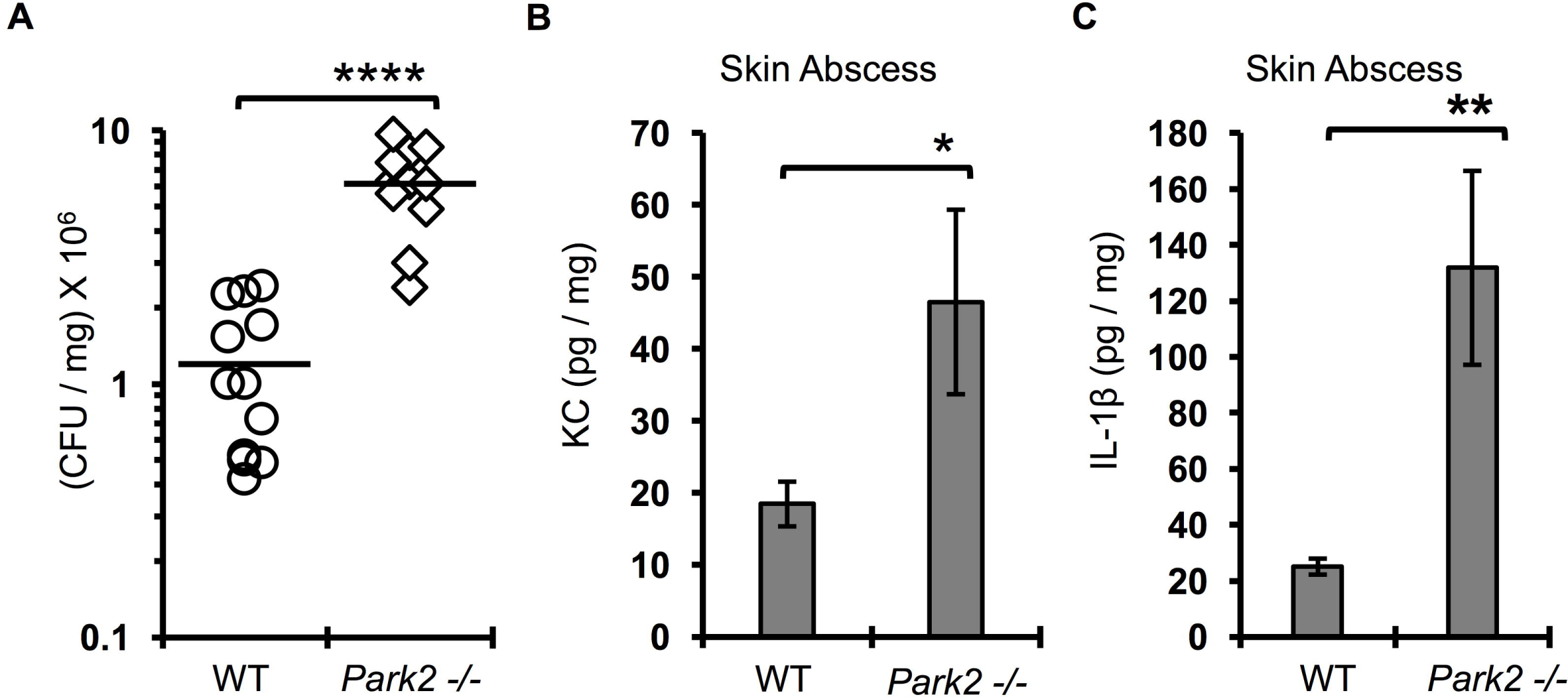
Parkin is essential for immunity during MRSA subcutaneous infection. (A) Bacterial burden in skin abscesses from male and female wild-type (WT) and Parkin deficient C57BL/6 mice (*Park2*^-/-^) infected subcutaneously with 10^7^ CFU of MRSA for 3 days. Horizontal lines represent the mean. Data are pooled from two independent experiments. (B-C) KC or IL1*β* cytokine levels in skin abscess homogenate of WT and *Park2*^-/-^ mice. Cytokine levels were quantified by ELISA. Graph bars represent mean of n=13 WT and 12 *Park2*^-/-^ mice pooled from 2 independent experiments. Error bars represent standard error of the mean (SEM). pValue: * < 0.05**, \***\* < 0.01 and **** < 0.0001.

### Inhibition of phagosomal acidification enhances MDV-mediated bactericidal capacity

Phagosomal acidification enhances macrophage antimicrobial function by reducing intracellular replication of certain pathogens such as *M. tuberculosis* (Sullivan et al., 2012). Conversely, other intracellular bacterial pathogens, such as *Salmonella enterica* serovar Typhimurium, are killed more efficiently when cells are treated with Bafilomycin A1 (Baf A1), an inhibitor of the vacuolar ATPase that prevents acidification and phagolysosomal fusion (Rathman et al., 1996). Notably, under stimulating conditions, Tom20^+^ MDVs accumulate when cells are treated with BafA1, presumably by preventing their resolution into the lysosomal network and subsequent degradation (Soubannier et al., 2012a; Sugiura et al., 2014). Therefore, we hypothesized that blockade of acidification and protein degradation in lysosomes with Baf A1 (Yoshimori et al., 1991) would increase MDV accumulation and thereby enhance bactericidal mROS delivery into MRSA-containing phagosomes. To determine if MDV accumulation enhanced macrophage bactericidal function, we first quantified MDV induction in infected macrophages treated with BafA1. We observed that BafA1 treatment increased Tom20^+^ MDV accumulation in MRSA-infected macrophages compared to DMSO-treated macrophages (Fig. 5A and 5B). BafA1-treated macrophages exhibited higher levels of mROS in MRSA-containing phagosomes and killed MRSA more efficiently compared to DMSO-treated cells (Fig. 5C-E). To further investigate how MDVs associate with MRSA-containing phagosomes, we performed transmission electron microscopy (TEM) on MRSA-infected macrophages treated with or without BafA1 at 4h pi. We could readily observe MRSA-containing phagosomes containing double membrane-bound vesicles (Fig. 5F). When cells were treated with BafA1, MRSA phagosomes appeared more spacious and the double membrane-bound vesicles could be easily observed inside the phagosome. To determine if these vesicles were derived from mitochondria, we performed immunogold labeling using anti-Tom20 antibody, followed by TEM. Although the immunogold fixation conditions decreased definition of phagosomal membranes, Tom20^+^ particles were observed in close proximity to the bacterial surface within the phagosomal space (Fig. S4). Collectively, these data suggest that preventing lysosomal acidification and protein degradation increased MDV accumulation and mH_2_O_2_ levels in the phagosome to augment MRSA killing.

**Figure 5.**
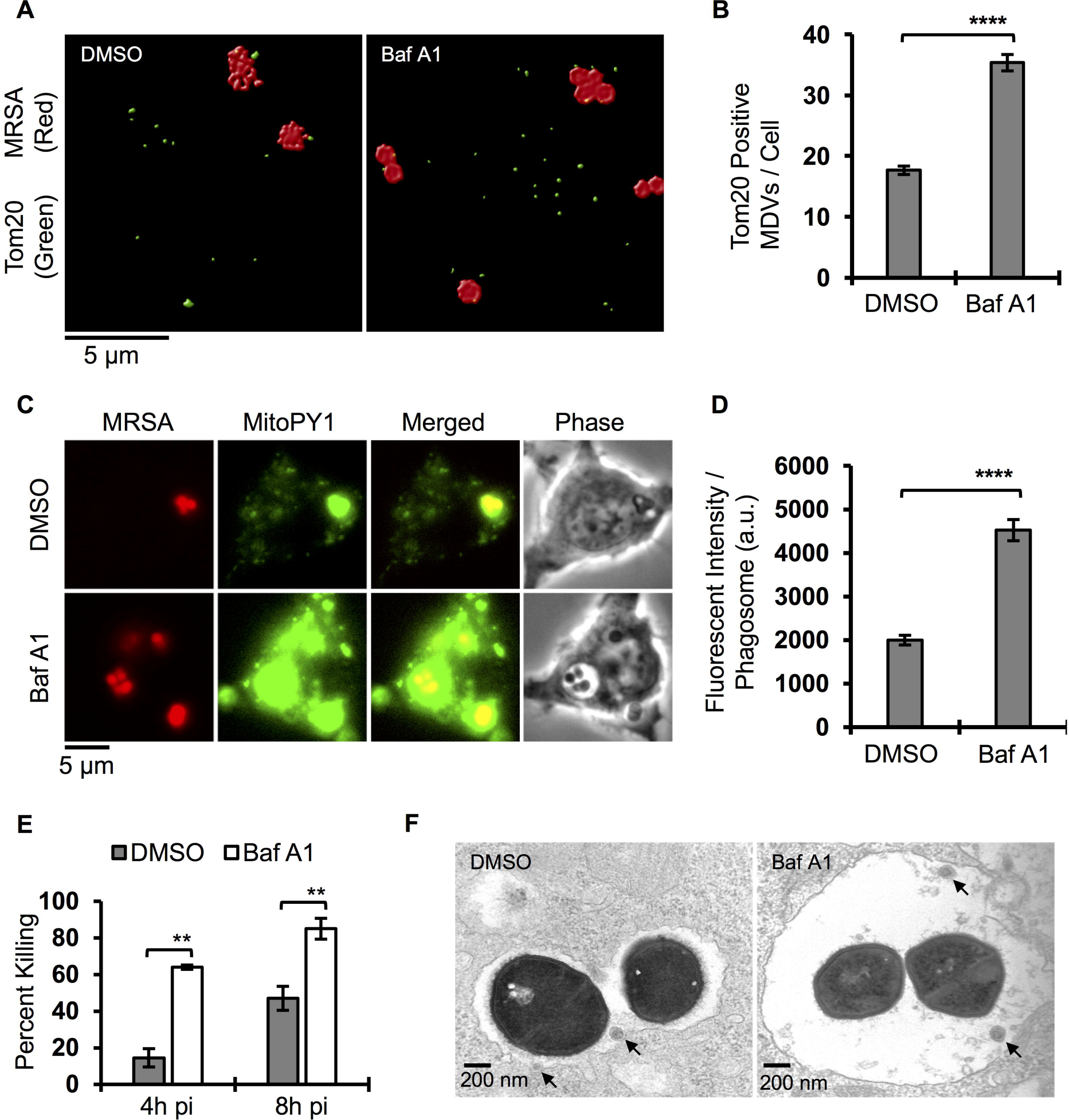
Blocked of phagolysosome fusion enhances MDV mediated mH_2_O_2_ killing. (A) Processed confocal microscopy representative images of RAW264.7 macrophages infected with MRSA-mCherry (red) for 4h and stained for Tom20 (green). Images were acquired by Leica TCS SP8 confocal scanning microscope and deconvoluted using Huygens essential software. Images were processed using the following setting; Threshold: 10%, Seed: 10% and Garbage: 50. Tom20 positive large objects (surface area > 0.4 µm^2^) were subtracted from the original images to define Tom20 positive small objects. (B) Quantification of Tom20 positive small objects (MDVs) per macrophage during MRSA-mCherry infection in the presence and absence of Bafilomycin A1. See detail in figure 3A. Data are presented as mean of at least 110 phagosomes from each condition pooled form three independent experiments +/− SEM. (C) Representative live wide-field microscopy images of RAW264.7 macrophages pulsed with MitoPY1 (green) and chased 4h post MRSA-mCherry (red) infection in the presence of Bafilomycin A1 (Baf A1, 100 nM) or controlled solvent (DMSO). (D) Mean fluorescent intensity (MFI) of MRSA-containing phagosomes when macrophages were being pulsed with MitoPY1 (green) and chased 4h post MRSA-mCherry (red) in the presence and absence of Bafilomycin A1 (Baf A1, 100 nM). Data represent mean of at least 321 phagosomes of each condition pooled from three independent experiments +/− SEM. (E) Percent of MRSA intracellular killing by RAW264.7 macrophages treated with Bafilomycin A1 (Baf A1, 100 nM) or control solvent (DMSO). Percent killing was quantified by the following formula [1 - (CFU _indicated time points_ / CFU_1h pi_)] × 100, which represent the percent difference in CFU obtained at indicated time point relative to 1h pi. Graph bars represent mean of n≥3 independent experiments +/− SD. (F) Transmission electron microscope images of macrophage MRSA-containing phagosomes. Macrophages are infected with MRSA for 4h in the presence and absence of Bafilomycin A1 (Baf A1, 100 nM). Images were acquired by using JEOL JEM-1400 Plus transmission electron microscope. Arrows indicate vesicles that are localized inside the phagosomes. pValue: **\***\* < 0.01 and **** < 0.0001.

### Sod2 is required for generation of bactericidal mH_2_O_2_ and is delivered to bacteria-containing phagosomes

Hydrogen peroxide is constitutively generated in the mitochondria from superoxide produced by electron transport that is converted by the mitochondrial manganese superoxide dismutase (Sod2) (Murphy, 2009). We therefore hypothesized that Sod2 was required for the bactericidal mH_2_O_2_ induced by MRSA infection. To define the spatial localization of Sod2 positive compartments relative to phagosomes, we performed confocal immunofluorescence microscopy on macrophages infected with MRSA or beads, staining with Sod2-and Lamp1-specific antibodies (Fig. 6A). Sod2 localized to the mitochondrial network in macrophages (Fig. S5) and large Sod2^+^ network objects were juxtaposed to both bead-and MRSA-containing phagosomes. However, small Sod2^+^ vesicles were induced during MRSA infection, and were present within MRSA-containing phagosomes. To further quantify Sod2^+^ vesicle accumulation in MRSA-containing phagosomes, we enumerated Sod2^+^ MDV located within a 1µm radius around bead-and MRSA-containing phagosomes, which were delineated by Lamp1^+^ staining, and found that MRSA infection increased Sod2^+^ MDV within phagosomes (Fig. 6B and 6C). To determine whether Sod2 was required for bactericidal activity, we stably knocked down Sod2 in RAW264.7 macrophages (Sod2 KD), which was confirmed by immunoblot analysis (Fig. 6D). We first tested the requirement of Sod2 in hydrogen peroxide and superoxide generation during MRSA infection (Fig. 6E, S6A and S6B). Compared to control cells, Sod2 KD cells failed to produce hydrogen peroxide upon MRSA infection while produced higher level of superoxide regardless of MRSA infection. Although Sod2 knockdown increased mitochondria superoxide production, it did not interfere with host cell death during MRSA infection (Fig. S6C). Importantly, Sod2 depletion impaired macrophage killing of MRSA compared to NT-control cells (Fig. 6F). Together, these results suggest that Sod2 enhances MRSA killing via generation of mH_2_O_2_, which is delivered by MDVs to bacteria-containing phagosomes.

**Figure 6.**
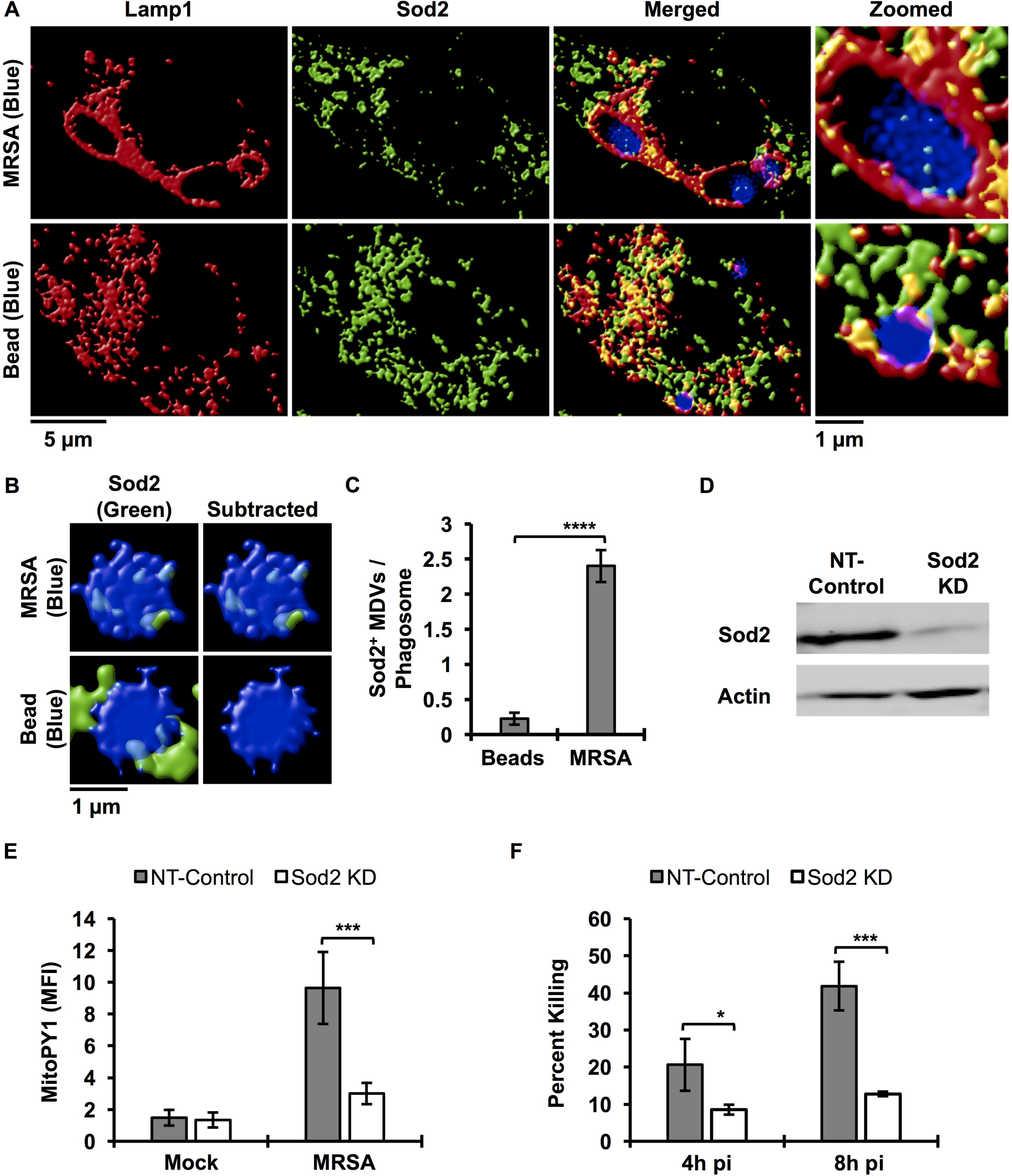
Sod2 is the mitochondria payload delivered to phagosome to promote mH_2_O_2_ killing. (A) Representative confocal microscopy images of RAW264.7 macrophages infected with MRSA (blue) or stimulated with bead (blue) for 4h and stained for Lamp1 (red) and Sod2 (green). Images were acquired using Leica TCS SP8 confocal scanning microscope and deconvoluted using Huygens essential software. (B) Confocal microscopy images of a magnified area of macrophages where bead or MRSA-containing phagosome is localized. Images in the right panel are the resulted images after subtracting large Sod2 positive objects (surface area > 0.4 µm^2^) from left panel images. (C) Quantification of Sod2 positive small objects (surface area < 0.4 µm^2^) of 2 µm^2^ area of the cell where beads or MRSA are localized. Large Sod2 positive objects were filtered out of the deconvoluted images and the number of remained Sod2 positive objects were enumerated per 2 µm^2^ areas of the cell where phagsosomes were localized. Data represent mean of at least 60 phagosomes from three independent experiments. (D) Immunoblots of cell lysate from RAW264.7 macrophages stably transduced with lentivirus-encoded shRNA for non-target (NT-Control) or Sod2 (Sod2 KD), probed with an anti-Sod2 antibody or anti-Actin antibody as a loading control. (E) Mean fluorescent intensity (MFI) of shRNA stably knockdown Sod2 (Sod2 KD) or non-target (NT-Control) macrophages after pulsed with MitoPY1 and chased 4h post MRSA infection. Samples were acquired by flow cytometry and analyzed by FlowJo software. MFI represent geometric mean of of n≥3 independent experiments +/− SD. (F) Intracellular MRSA killing by Sod2 knockdown (Sod2 KD) and non-target (NT-Control) macrophages was quantified by using the following formula [1 - (CFU _indicated time_ points / CFU_1h pi_)] and expressed as percentage. Graph bars represent mean of percent killing of n≥3 independent experiments +/− SD. pValue: * < 0.05, **\***\*\* < 0.001 and **** < 0.0001.

## DISCUSSION

The generation of anti-microbial reactive oxygen intermediates is an integral weapon in the innate immune arsenal. We find that infection by MRSA stimulates the production of mitochondrial ROS, specifically mitochondrial hydrogen peroxide. Our results identify Sod2 as a key enzyme responsible for infection-induced mH_2_O_2_ generation. Moreover, these studies reveal delivery of the Sod2/mH_2_O_2_ payload to the phagosome by mitochondria-derived vesicles as a novel mechanism for anti-microbial killing. Stimulation of mH_2_O_2_-containing MDV during infection required TLR signaling and employed a Parkin-dependent pathway. Taken together, our data support a model where Sod2-driven mH_2_O_2_ production and accumulation via MDV delivery establish a potent killing ground for bacterial pathogens within the macrophage phagosome.

Generation of MDV has been described as a quality control mechanism that could transport damaged respiratory chain complexes from mitochondria to the endolysosomal compartment for degradation (Soubannier et al., 2012a). More recent studies reveal that MDV can carry out diverse functions within the cell, and likely represent multiple vesicle populations with distinct cargo related to their function. Sugira, *et al*, provided evidence for MDV generation as a critical step in peroxisome biogenesis, where MDV containing the peroxisomal proteins, Pex3 and Pex14, fuse with ER-derived vesicles, containing Pex16, resulting in import-competent organelles (Sugiura et al., 2017). This model of peroxisome biogenesis adds a new dimension to our understanding of how MDVs carry out mitochondrial communication with other organelles. Our work further identifies the phagosome as a new destination for MDVs that is revealed by infection.

Soubannier *et al* showed that different proteins are incorporated in MDVs when mitochondria are stressed with different stimuli, implying the existence of mechanisms to package and transport specific cargos. For instance, exogenous ROS applied to isolated mitochondria results in MDVs enriched in the outer membrane protein, VDAC. However, when ROS is generated by isolated mitochondria via inhibition of complex III by Antimycin A, MDVs carried complex III subunit core2 without enrichment of VDAC (Soubannier et al., 2012b). A recent study showed that in response to heat shock or LPS, MDVs delivered mitochondrial antigens to antigen-loading compartment independently of Parkin (Matheoud et al., 2016). In contrast, when cells are treated with Antimycin A to stimulate mitochondrial ROS production, MDVs are targeted to lysosomes in a Parkin-dependent manner (McLelland et al., 2014). In the context of infection, our data demonstrated that MDVs are generated in a Parkin-dependent manner, similar to MDVs destined for lysosomes. Moreover, we showed that infection-induced MDVs contain mitochondria-specific enzyme, Sod2. Collectively, our data and others support the idea that specific stimuli define the packaging and destination of MDV subsets.

In the context of the mitochondrial matrix, Sod2 cooperates with other enzymes and anti-oxidant proteins to detoxify superoxide generated by oxidative metabolism. Notably, dysregulation of Sod2 is associated with many human diseases, some of which are associated with increased inflammation (Flynn and Melov, 2013). Altering the spatial distribution of Sod2 could disrupt the coordinated ROS detoxification process, resulting in increased hydrogen peroxide production without the capacity to further reduce to water. Previous studies provide some evidence to support this hypothesis. Overexpression of Sod2 increased the steady state of hydrogen peroxide in cancer cells, and may contribute to tumor invasion and metastasis (Nelson et al., 2003; Ranganathan et al., 2001). In contrast, decreasing Sod2 activity in Sod2-heterozygous mice led to higher production of superoxide radical when measured by aconitase activity, which correlate with an increase in mitochondrial oxidative damage (Williams et al., 1998). Consistent with these data, our results showed that decreasing macrophage Sod2 leads to elevation of steady state levels of mitochondrial superoxide and decreased generation of hydrogen peroxide during MRSA infection. As a consequence, Sod2-deficient macrophages (Sod2 KD) failed to kill MRSA. Although we were not able to successfully generate live Sod2-deficient mice, studies of Sod2 knockdown in zebrafish demonstrate a protective role for Sod2 during infection by *Pseudomonas aeruginosa* (Peterman et al., 2015). Our studies have identified MDVs as a new delivery mechanism by which antimicrobial effectors can be trafficked into the macrophage phagosome. We propose that removal of a subset of Sod2 protein from the coordinated redox environment of the mitochondrial network through packaging into MDVs allows repurposing of this enzyme from detoxification to anti-microbial defense.

## Acknowledgments

This work was supported by NIH award R21 AI101777 (M.X.O). B.H.A. was supported by NIH T32 AI007538, T32 HLL007517 and an American Heart Association Post-doctoral fellowship. We thank O’Riordan lab members for many helpful discussions. We gratefully acknowledge the Center for Live Cell Imaging (CLCI), Microscopy and Image Analysis Laboratory (MIL) and Cancer Center Immunology Core at the University of Michigan Medical School. We thank Dr. T. Merkel (FDA) for *Tlr*2/4/9^-/-^ femurs.

## Author Contributions

B.A. performed the experiments; B.A. and M.O. designed the experiments and wrote the manuscript; T.S. assisted in experimental preparation.

## STAR Methods

### CONTACT FOR REAGENT AND RESOURCE SHARING

Reagents and resources can be obtained by directing requests to the Lead Contact, Mary O’Riordan

### EXPERIMENTAL MODEL AND SUBJECT DETAILS

#### Mice

Wild-type C57BL/6 and *Park2*^-/-^ are purchased from the Jackson Laboratory. *Tlr2/4/9*^-/-^ have been described previously (Abuaita et al. 2015). All mice were maintained according to an approved protocol in the Unit for Laboratory Animal Medicine (ULAM) facilities at the University of Michigan Medical School.

#### Cells

Primary bone marrow derived macrophages (BMDMs) were prepared by flushing mouse femurs in DMEM supplemented with 100 units/ml of Pen/Strep. Cells were differentiated by incubation in BMDM medium (50% DMEM, 2 mM L-glutamine, 1 mM sodium pyruvate, 30% L929-conditioned medium, 20% heat-inactivated fetal bovine serum (FBS), 55 µM 2-mercaptoethanol, and Pen/Strep). L-929 and HEK 293T cells were cultured in MEM supplemented with 2 mM L-glutamine, 1 mM sodium pyruvate, 1mM Non-essential amino acid (NEAA), 10 mM HEPES, and 10% heat-inactivated FBS). RAW 264.7 cells were cultured in RPMI 1640 containing 2 mM L-glutamine and 10% heat-inactivated FBS. All cells were incubated at 37°C in 5% CO_2_.

#### Bacterial infections

USA300 LAC, a community associated methicillin-resistant *Staphylococcus aureus* strain (MRSA) and its isogenic strain harboring p*SarA*-mCherry plasmid (MRSA-mCherry) (Boles and Horswill, 2008), were maintained at −80°C in LB medium containing 20% glycerol. All strains were cultured in tryptic soy agar (TSA, Becton Dickinson), and selected colonies were grown overnight at 37°C with shaking (240 rpm) in liquid tryptic soy broth. Bacteria were pelleted, washed and re-suspended in PBS. The bacterial inoculum was estimated based on OD_600_, and verified by plating serial dilutions on TSA plates to determine colony forming units (CFU). Macrophages were infected at a multiplicity of infection (MOI) of 20 in culture medium without antibiotic for 45 minutes. Infected macrophages were washed three times with PBS and incubated in medium containing 100 µg/ml of gentamicin to kill extracellular bacteria for 15 minutes. Media was exchanged with media containing 50 µg/ml of gentamicin for the remaining time of the experiments.

### METHOD DETAILS

#### Macrophage bactericidal activity

Macrophages were seeded in a 24-well tissue culture treated plate at a density of 1.5 × 10^5^ cell/well. The next day, macrophages were infected with MRSA (MOI of 20). The number of intracellular bacteria was determined by washing infected macrophages with PBS, lysing with 0.1% NP-40, and enumerating bacterial CFUs via serial dilution on agar plates. The percentage of killed MRSA was calculated by the following formula [1 - (CFU _indicated time points_ / CFU_1h pi_)] × 100, which represents the percent difference between CFU at indicated time points relative to 1h pi. Where indicated, macrophages were pre-incubated with NecroX-5 (10 µM), Bafilomycin A1 (100 nM) or Mdivi-1 (25 µM) for 30 minutes before infection, and all inhibitors were maintained throughout the experiment.

#### Mouse infection

Subcutaneous MRSA infection was performed as previously described (Tseng et al., 2011). Male and Female C57BL/6 mice and *Park2*^*-/-*^ were shaved on the right flank. Mice were inoculated with 10^7^ bacteria in 100 µl of PBS subcutaneously on the shaved area of the skin using a 27 gauge needle. Mice were sacrificed on day 3 post-infection and skin abscesses were excised, weighed and homogenized in PBS. Total CFU per mouse abscess were enumerated by serial dilution and plating on TSA agar. Total CFU was converted to CFU/mg of tissue weight. Cytokines were quantified by ELISA at the University of Michigan ELISA core and converted to pg/mg of tissue weight.

#### Cellular mROS measurement

Macrophages were plated in 60 mm non-treated dishes and treated with 10 µM MitoPY1 (TOCRIS) or 10 µM MitoSOX (Life Technology) for 1 hour. Macrophages were washed three times with media and only when indicated, macrophages were treated for 30 minutes with NecroX-5 (10 µM) or control solvent prior to infection with MRSA (MOI of 20). For positive controls, macrophages were treated with 100 µM hydrogen peroxide or 10 µM Antimycin A for 1 hour to induce the oxidation of MitoPY1 or MitoSOX, respectively. Macrophages were subjected to flow cytometry and data were analyzed with FlowJo software. The mean fluorescence intensity for each condition was determined as the geometric mean.

#### Phagosomal mROS measurement

Macrophages were plated in 35 mm glass bottom dishes (MatTek). The next day, macrophages were treated with MitoPY1 (10 µM) for 1 hour, washed three times with media and stimulated with inert red fluorescent beads or infected with live or dead (inactivated by paraformaldehyde) red fluorescent MRSA harboring p*SarA*-mCherry (MOI of 20). Macrophages were imaged in Ringer buffer (155 mM NaCl, 5 mM KCl, 1 mM MgCl2.6H2O, 2 mM NaH2PO4.H2O, 10 mM HEPES, and 10 mM Glucose) with an Olympus IX70 inverted live-cell fluorescence microscope. Fluorescence images were further processed by MetaMorph imaging software. For quantification of mROS association with phagosomes, the mean fluorescence intensity of MitoPY1 at the phagosome area was measured by ImageJ. Phagosomal regions in the cell images were defined by the location of red fluorescent beads or bacteria. For BMDM experiments, a ratiometric mean fluorescence intensity of the phagosomal area over the mean fluorescence intensity of the cell was calculated. This was done because Park2 deficient macrophages have higher global MitoPY1 fluorescence intensity when compared to WT macrophages prior to infection.

#### Confocal microscopy

Macrophages were seeded onto microscope cover glass and infected with MRSA (MOI of 20) or stimulated with fluorescent beads. Cells were fixed at 4h pi with 3.7% paraformaldehyde at room temperature for 20 minutes and permeabilized with PBS contain 0.1% Triton X-100 for 15 minutes. MRSA were stained using chicken antiprotein A antibody conjugated to biotin (Abcam ab18598) in staining buffer (PBS, 0.1% Triton X-100, 5% BSA, and 10% normal goat serum). Host proteins were stained using mouse anti-Sod2 (Abcam ab110300, clone 9E2BD2), Rat-anti-Lamp1 (DSHB, clone 1D4B), and anti-Tom20 (Santa cruz, FL145). Secondary antibodies (goat anti-mouse (Alexa-488), goat anti-Rat (Alexa-594) and Streptavidin (Alexa-405) were used according to manufacturer’s procedure. Cover glasses were mounted on microscope slides using Prolong Diamond (Life Technology). Cells were imaged using a Leica TCS SP8 confocal microscope and deconvoluted using Huygens essential software by scientific volume imaging using the following criteria; Threshold (10%), Seed (10%) and Garbage Volume (50). To define MDVs, large objects (surface area larger than 0.4 µm^2^) were filtered out from the mitochondrial fluorescence labeled channel and the number of remaining objects per cell were recorded.

#### Transmission electron microscopy

Macrophages were infected with MRSA (MOI of 20) in the presence of Bafilomycin A1 (100 nM) or control DMSO. Infected macrophages were fixed at 4h pi with 2.5% glutaraldehyde for at least 1h at room temperature, then overnight at 4°C. For immuno-gold staining, infected macrophages were stained for Tom20 prior to fixation with glutaraldehyde according to manufacturer’s procedure (AURION). Briefly, cells were fixed with 3.7% paraformaldehyde at room temperature for 20 minutes and permeabilized with Sorenson’s buffer containing 0.1% Triton X-100 for 15 minutes. Cells were blocked with the AURION blocking solution (AURION-BSA-c) and stained using primary anti-Tom20 antibody (Santa Cruz, FL145) and secondary goat anti-rabbit ultra-small gold antibody (AURION). Silver stain enhancement was carried out by using the AURION R-GENT SE-EM according to reagent protocol (AURION). Glutaraldehyde fixed samples were washed with Sorenson’s buffer 3-times before post-fixing in 2% osmium tetroxide in Sorenson’s buffer for 1h at room temperature. Samples were washed again 3-times with Sorenson’s buffer, then dehydrated through ascending concentrations of ethanol, treated with propylene oxide, and embedded in EMbed 812 epoxy resin. Semi-thin sections were stained with toluidine blue for tissue identification. Selected regions of interest were ultra-thin sectioned to 70 nm and post stained with uranyl acetate and Reynolds lead citrate. Sections were examined using a JEOL JEM-1400 Plus transmission electron microscope (TEM) at 80 kV.

#### Generation of RAW264.7*ΔIre1-α* and RAW264.7 shRNA stable knockdown cells

The generation of lentivirus for CRISPR-Cas9 knockout and shRNA knockdown was done by using HEK293T packaging cells, which were grown in DMEM with 10% FBS. The virus particles were produced by transfecting the cells with the TRC shRNA encoded plasmid (pLKO.1) or guided RNA (gRNA) encoded plasmid (lentiCRISPRv2) along with the packaging plasmids (pHCMV-G, and pHCMV-HIV-1) (Kulpa et al., 2013) using FUGENE-HD transfection reagent (Promega). Media was changed after 24h and virus particles were collected after 72h post-transfection. A total of 2 ml of medium containing virus were concentrated ten-fold by ultracentrifugation at 24,000 rpm for 2h at 4°C and used to transduce RAW264.7 cells. Transduced cells were selected with puromycin (3 μg/ml). The mouse Sod2 specific shRNA plasmid with the sense sequence of (GCTTACTACCTTCAGTATAAA) and the non-target control shRNA plasmid were purchased from Sigma-Aldrich. The efficiency of knockdown was monitored by immunoblot analysis using anti-Sod2 antibody (Santa Cruz). Anti-Actin antibody was used as a loading control (Fisher Scientific). The mouse IRE1-*α* specific gRNA sequence of (CTTGTTGTTTGTCTCGACCC) and the non-target gRNA control sequence of (TCCTGCGCGATGACCGTCGG) were cloned into lentiCRISPRv2 according to the Feng Zhang lab protocol (Sanjana et al., 2014). Single clones of RAW264.7*ΔIre1-α* were isolated and confirmed by the absence of IRE1-*α* protein by immunoblot using anti-IRE1-*α* antibody (clone 14C10, Cell Signaling). Anti-GAPDH antibody was used as a loading control (Santa Cruz). RAW264.7*ΔIre1-α* clone was also confirmed by absence of its endonuclease activity when cells were treated with endoplasmic reticulum stress inducer Thapsigargin (5 µM) by *xbp1* splicing assay (Figure S1) as previously described (Abuaita et al., 2015).

#### Cell death assay

Cell death was measured by flow cytometry using SYTOX green dead cell stain according to manufacturer’s protocol (Life Technology). Briefly, macrophages were incubated with SYTOX green dead cell stain (30 nM) in HBSS for 20 minutes at room temperature prior to flow cytometry analysis using 488 excitation and 530/30 emission. Digitonin (0.01%, Sigma Aldrich) was used as a positive control to permeabilize the plasma membrane. The percent of SYTOX positive cells was determined by gating against mock unstained cells.

### QUANTIFICATION AND STATISTICAL ANALYSIS

Data were analyzed using Excel 2016 and Student’s unpaired two-tailed t-test was applied. The mean of at least three independent experiments was presented with error bars showing standard deviation (SD) or standard error of the mean (SEM), which is indicated in figure legends. *P* values of less than 0.05 were considered significant and designated by: \**P* < 0.05, \*\**P* < 0.01, \*\*\**P* < 0.001 and \*\*\*\**P* < 0.0001. All statistically significant comparisons within experimental groups are marked.

### DATA AND SOFTWARE AVAILABILITY

RAW data are available upon request, which should be directed to the Lead Contact. There was no proprietary software used in this study.

### ADDITIONAL RESOURCES

mCherry-Mito-7 plasmid was used under material transfer agreement (MTA).

## Supplemental Information

**Figure S1. IRE1α endonuclease activity is absent in RAW264.7ΔIRE1α macrophage cell line in response to ER Stressor, Thapsigargin**

RT-PCR analysis of *Xbp1* mRNA splicing in single clone of RAW 264.7 macrophages stably transduced with IRE1*α* specific guided RNA (gRNA) or non-target (NT-control). Macrophages were treated with 5 µM thapsigargin (TG) to induce IRE1*α* endonuclease activity or control DMSO. RT-PCR products were digested with *PstI* endonuclease. Because unspliced *Xbp1* mRNA contains a *PstI* site within the 26 spliced region, the digested RT-PCR products yield two smaller fragments representing the unspliced (U) *Xbp1* and one larger fragment representing the spliced (S) *Xbp1*. RT-PCR image is representative of n≥3 independent experiments.

**Figure S2. TLR2/4/9 signaling is dispensable for MRSA-induced mH_2_O_2_ but is essential for macrophage bactericidal activity**

(A) Mean fluorescent intensity (MFI) of wild-type (WT) and *Tlr2/4/9* triple knockout (TLR2/4/9 KO) macrophages that were loaded with MitoPY1 and infected with MRSA. Macrophages were subjected to flow cytometry at 4h pi and collected data were analyzed by FlowJo software for geometric mean.

(B) Percent of MRSA intracellular killing by WT and TLR2/4/9 KO macrophages.

Macrophage killing efficiency was calculated by using the following formula [1 -(CFU_indicated time points_ / CFU_1h pi_)] × 100, which represent the percent difference in CFU obtained at indicated time point relative to 1h pi. Data are expressed as percentage.

Graph bars represent mean of of n≥3 independent experiments +/− SD. pValue: **\***\* < 0.01 and **** < 0.0001.

**Figure S3. Tom20 positive MDVs are defined by small objects that are stained with Tom20**

(A) Representative confocal images of MRSA infected and beads internalized macrophages that stained for Tom20. Images were acquired by Leica TCS SP8 confocal scanning microscope and deconvoluted using Huygens essential software. Images were processed using the following setting; threshold: 10%, Seed: 10% and Garbage: 50. Right panel, Tom20 positive large objects (surface area > 0.4 µm^2^) were subtracted from left panel Images.

(B) Quantification the number of Tom20 positive objects per beads internalized or MRSA infected macrophage that fall into indicated range of surface size bins.

Discontinuous line was drawn to point out the surface area size bin smaller than 0.4 µm^2^, which was chosen to quantify Tom20 small objects (MDVs).

**Figure S4. Tom20 positive vesicles are localized in the lumen of MRSA phagosomes**

Transmission electron microscopy representative mages of MRSA infected macrophages when treated with Bafilomycin A1 (Baf A1, 100 nM) or control DMSO. Tom20 was stained with immune-gold particles followed by silver enhancement. Left panel; images showing whole cell, middle panel; magnified images showing mitochondria, right panel; images showing MRSA phagosome. Arrows were drawn to indicate Tom20 positive gold particles present in MRSA phagosome lumen.

**Figure S5. Macrophage Sod2 is localized with Tom20**

Representative confocal microscopy images of RAW264.7 macrophages were stained with Sod2 and Tom20. Images were acquired with Leica TCS SP8 confocal scanning microscope and deconvoluted using Huygens essential software. Pearson Correlation Coefficient (PCC) was determine on the deconvoluted images using Huygens essential software and presented as mean of at least 230 cells +/− SD pooled from three different experiments.

**Figure S6. Knockdown macrophage Sod2 increases mitochondria superoxide production without interfering with host cell death during MRSA infection**

(A) Representative histogram plots are shown when macrophages were pulsed with MitoSOX for 1h and chased 4h after stimulated with Antamycin A (10 µM), infected with MRSA at MOI of 20 or left untreated (Mock). Percent of MitoSOX positive cells was determined by gating against unstained cells. Percent of MitoSOX high cells was determined by gating against the MitoSOX first peak.

(B) Quantification of the percentage MitoSOX high from panal A. Data represent the percentage of cells with MitoSOX high relative to total MitoSOX positive cells.

(C) Percent of live macrophages was determined by Sytox green dead cell staining. NT-control and Sod2 KD macrophages were infected with MRSA for 4h (MOI of 20), treated with 0.01% Digitonin (Digitonin) to induce cell death or left untreated (Mock). Cells were stained with 30 nM of Sytox green in HBSS buffer and subjected to flow cytometry. Data are analyzed by FlowJo software. Percent of Sytox green positive cells were determined by gating against live cell peak. Graph bars represent mean of n≥3 independent experiments +/− SD. pValue: * < 0.05 and **\***\* < 0.01.

**Movie S1. mH_2_O_2_ accumulates in MRSA-containing phagosome**

Time-lapse movie of RAW264.7 macrophage after being pulsed with mitochondria-targeted H_2_O_2_ fluorescent sensor, MitoPY1 (green fluorescent) for 1h and infected with MRSA-mCherry (Red fluorescent). Images were acquired using inverted IX-70 Olympus live fluorescent microscope and analyzed using MetaMorph imaging software. Time clock Minutes : Seconds.

